# Statistically correcting dynamical electron scattering improves refinement of protein nanocrystals, including charge refinement of coordinated metals

**DOI:** 10.1101/2020.07.08.191049

**Authors:** Thorsten B. Blum, Dominique Housset, Max T.B. Clabbers, Eric van Genderen, Maria Bacia-Verloop, Ulrich Zander, Andrew A. McCarthy, Guy Schoehn, Wai Li Ling, Jan Pieter Abrahams

## Abstract

Electron diffraction allows protein structure determination when only nanosized crystals are available. Nevertheless, multiple elastic (or dynamical) scattering, prominent in electron diffraction, is a concern. Current methods for modeling dynamical scattering by multi-slice or Bloch wave approaches are not suitable for protein crystals because they are not designed to cope with large molecules. Here, we limited dynamical scattering of nanocrystals of insulin, thermolysin, and thaumatin by collecting data from thin crystals. To accurately measure the weak diffraction signal from the few unit cells in the thin crystals, we used a low-noise hybrid-pixel Timepix electron counting detector. The remaining dynamical component was further reduced in refinement using a likelihood-based correction, which we introduced previously for analyzing electron diffraction data of small molecule nanocrystals and adapted here for protein crystals. We show that the procedure notably improved the structural refinement, allowing in one case the location of solvent molecules. It also allowed the refinement of the charge state of bound metal atoms, an important element in protein functions, through B-factor analysis of the metal atoms and their ligands. Our results clearly increase the value of macromolecular electron crystallography as a complementary structural biology technique.

## Introduction

The strong interaction of electrons with matter favors electron crystallography for solving three-dimensional (3D) structures from beam-sensitive sub-micron-sized crystals, such as protein nanocrystals (Clabbers and Abrahams, 2018; Henderson, 1995). With cryo-electron microscopy, protein crystals can be vitrified and studied in their native hydrated states. In contrast to real-space techniques like single particle analysis or electron tomography, electron diffraction data are recorded in the diffraction plane. The major advantage of image-based methods over diffraction is that they provide experimental data with crystallographic phases. However, compared to diffraction methods, this advantage comes at a severe price in terms of signal strength, as an electron dose required for a single high-resolution still image is sufficient for a full 3D diffraction rotation data set of the same crystal (Clabbers and Abrahams, 2018). Indeed, various protein structures have been solved with diffraction data collected with continuous rotation of 3D crystals recently (Clabbers et al., 2017; de la Cruz et al., 2017, Nannenga et al., 2014; Shi et al., 2013; Xu et al., 2018; Yonekura et al., 2015).

Early attempts to use electron diffraction in structural biology were promising but were mainly limited by the electron detectors. Coupled with phase information obtained from images, structures were solved to 6-7 Å in some early studies (Subramaniam et al., 1997; Unwin and Henderson, 1975). Only a few still diffraction patterns could be recorded from any single (2D) protein crystal with the imaging plates or charge coupled detectors used in these studies. The limitation is practically due to the significant radiation damage imposed on the crystals necessary to obtain sufficient signal strength on these low-sensitivity detectors. Recent development of direct electron detectors has drastically reduced the required dose to achieve data quality suitable for data analysis.

In parallel, another important technological advancement was achieved in CMOS and hybrid pixel detectors that allow low-noise and high-speed imaging at high dynamic ranges, which is especially important for diffraction studies. Due to the very low curvature of the Ewald sphere for diffraction in typical transmission electron microscopes operating in the 100 – 300 kV range, still diffraction patterns yield only quasi 2D planes in diffraction space. Fast detectors allow the 3D sampling of the reciprocal space by collecting diffraction data from continuously rotating samples and the straightforward integration of the reflections fully recorded during the rotation (Gemmi et al., 2019; van Genderen et al., 2016).

Electron diffraction data collection with fast sensitive electron detectors on continuously rotating crystals has yielded data with a quality and quantity that permits standard X-ray crystallography programs such as MOSFLM (Leslie, 2006), XDS (Kabsch, 2010) or Dials (Clabbers et al., 2018) to process the electron diffraction data. Data integration became straightforward using a range of mature public domain software packages, each with a wide user base and supported by longstanding crystallography software consortia. There is also a range of standard crystallographic software for subsequent structure solution by molecular replacement or direct methods (if permitted by the resolution) and refinement, e.g. CCP4 suite (Winn et al., 2011) or SHELX (Usón and Sheldrick, 1999).

Nevertheless, such programs do not consider multiple elastic (dynamical) scattering. Dynamical scattering is negligible in X-ray diffraction but not so in electron diffraction due to the strong interaction of electrons with matter. As multiple elastic scattering coincides with kinematic Bragg angles, the measured intensities can no longer be simply approximated as the square of the structure factors. Dynamical diffraction theory has been successfully used for dynamical structure refinement for electron diffraction data (Palatinus et al., 2017). However, such methods require the knowledge of the atomic structure, the shape, and the orientation of the crystal for the modeling of the electron wavefunction travelling through the crystal. Currently, such an approach is only feasible for small molecules with small unit cells.

Data collection strategies may be used to help minimize multi-scattering events for macromolecule crystals. In particular, thin crystals can be selected to reduce the scattering path length. A data collection protocol, in which the crystal is continuously rotated about a random axis, will reduce dynamical scattering through integration (and thus averaging) of Bragg spots measured in different crystal orientations (Clabbers and Abrahams, 2018; Subramanian et al., 2015). The remaining dynamical scattering can be further corrected using a likelihood-based method described by Clabbers et al. (Clabbers et al., 2019). As dynamical scattering increases the intensity of weak reflections and decreases the intensity of strong reflections, weak diffraction data are, on average, overestimated. Applying a statistical correction to reduce overestimated weak intensities as a function of resolution can therefore significantly improve structure factor accuracy.

Another feature that distinguishes electron diffraction from other diffraction techniques is that electrons are scattered by the Coulomb potential in the crystal. Atomic scattering factors for electrons therefore depend strongly on the charge state of the atoms at low and medium resolution (Yonekura and Maki-Yonekura, 2016). This feature of electron diffraction offers a unique opportunity in structural biology as charged groups are central to protein functions such as enzyme catalysis (Warshel et al., 2006). Metal ions, in particular, often act as catalytic or structural cofactors in enzymes that participate in important metabolic pathways. Identifying partial charge states of metals in proteins will be crucial in understanding the reactivity of functional proteins. Refinement against electron diffraction data with different metal ion charge states will allow us to detect their partial charges in the protein crystals (Yonekura and Maki-Yonekura, 2016).

Here, we collected continuous rotation electron diffraction data of protein nanocrystals of bovine insulin, thermolysin from *Bacillus thermoproteolyticus*, and thaumatin from *Thaumatococcus daniellii*. Thin crystals were chosen for data collection using the Timepix hybrid pixel detector. To further reduce the dynamical scattering component of the data, we applied the likelihood-based approach to correct for the systematic overestimation of weak electron diffraction intensities. Here, we improve the method of Clabbers and collaborators (Clabbers et al., 2019) by including all measured intensities, including negative ones, in the calculation of correction parameters. We show that this correction notably improves crystallographic refinement. This improvement allowed exploring the scattering factors for various charge states of coordinated metal ions in insulin and thermolysin for the final refinement. For insulin, we also compared the electron diffraction results with X-ray diffraction results obtained from micron-sized crystals obtained from the same preparation.

## Results

### Overall description of the structures

The data processing and refinement statistics for the three proteins are shown in table 1. The structure of bovine insulin was determined by both X-ray and electron diffraction using crystals from the same batch, *i*.*e*. grown in the exact same condition. The X-ray structure was refined at 2.3 Å resolution, sufficient to yield a reference structure for the electron diffraction structure. The structure is very close to the T6 bovine insulin structure (2A3G protein data bank (PDB) entry) (r.m.s. difference of 0.34 Å on Cα atoms) and belong to the same space group with similar unit cell dimensions, albeit obtained from crystals grown in different crystallization conditions. The asymmetric unit contains two insulin molecules as well as two Zn ions and two Cl ions located on the crystallographic 3-fold axis, and 23 water molecules. The electron density of these insulin crystals is well defined for all amino acids except for residue 30 of chain B and residues 1 and 2 of chain D.

**Table 1:**
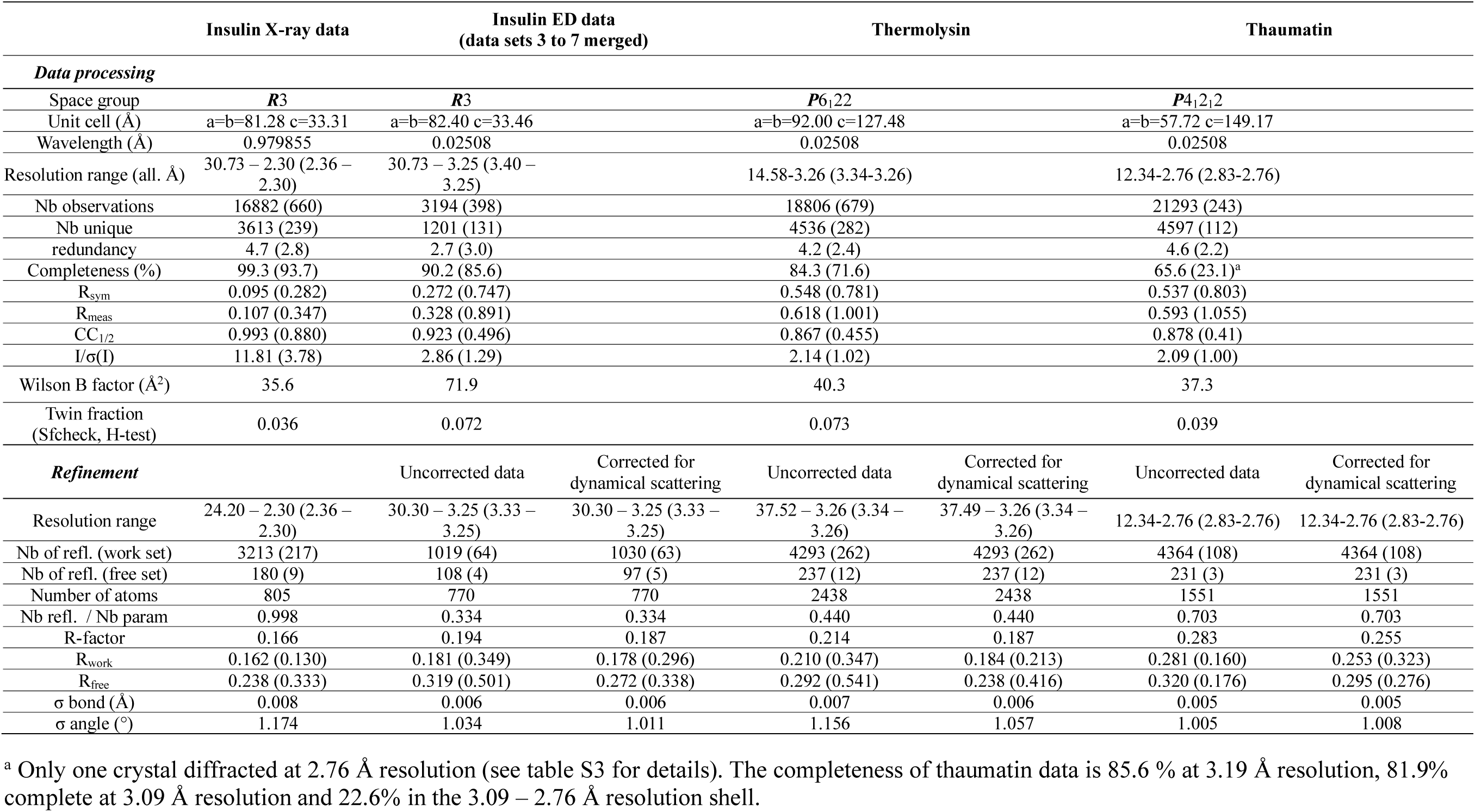
Data processing and refinement statistics for X-ray diffraction data of insulin and electron diffraction (ED) data of insulin, thermolysin, and thaumatin.

The insulin structure refined against electron diffraction data at 3.25 Å resolution agrees well with the X-ray structure refined at 2.3 Å resolution in this study (r.m.s. difference of 0.38 Å on Cα atoms). Similar to the electron density map from X-ray, the Coulomb potential map is well defined along all the polypeptide chains, except for residues 29-30 of chain B and residues 1 and 2 of chain D. However, no solvent molecules could be identified in the Coulomb potential map except for one Zn ion, which is clearly visible in the map and corresponds to the one chelated by histidine 10 of chain D (and its two symmetry mates). The Zn site on the B chain is vacant. This observation is consistent with the fact that the insulin crystals were diluted and crushed in pure water, which might have caused Zn ions to diffuse out of the nanocrystals during the vortexing step (see Materials and Methods). Moreover, the overall B-factor estimated from the Wilson plot is higher for the electron diffraction data (71.9 Å^2^) than for the X-ray data (35.6 Å^2^), suggesting that the crushing step and/or the loss of Zn ions may have introduced some disorder in the insulin crystals.

The thermolysin structure refined against electron diffraction data at 3.26 Å resolution also exhibits a very good fit with the Coulomb potential map for all the 316 residues. One Zn ion and 4 Ca ions could clearly be observed in the map and were included in the refined model. The final model is very similar to the X-ray crystallographic structure of the same protein at 2.0 Å resolution (1FJ3) with an r.m.s. difference of 0.49 Å for 314 Cα atom pairs.

The thaumatin structure was refined against electron diffraction data up to 2.76 Å resolution. The fit with the Coulomb potential map is good for all the 207 residues. The structure is close to the 6C5Y PDB entry, an X-ray crystallographic structure refined at 2.5 Å resolution obtained from microcrystals (r.m.s. difference of 0.42 Å for 202 Cα atom pairs). No solvent molecules were identified except for one putative Cl ion on a twofold axis and coordinated by the guanidinium group of Arg82.

Whereas the refinement statistics given in table 1 are far from ideal compared to X-ray crystallography standards, they are in line with previous structures refined against electron diffraction data at similar resolution. Indeed, previous protein structures solved by 3D electron diffraction deposited in the Protein Data Bank in the 2.7–3.4 Å resolution range exhibit R_work_ and R_free_ in the range of 0.21-0.32 and 0.25-0.34, respectively. The gap between R_work_ and R_free_ increases as expected when the data-to-parameter ratio decreases as observed in the different data sets here. In general, refinement against X-ray diffraction data yields better values for R_work_ than refinement against electron diffraction data. Undoubtedly, multiple scattering and lower I/σ(I) ratios contributed to this discrepancy. The relatively large gap observed for the electron diffraction data of insulin is likely to be due to both the low resolution (3.25 Å) and the low solvent content (∼35%) in the crystals. The rather low overall completeness of thaumatin data comes from the fact that only one crystal diffracted up to 2.76 Å resolution, while the others diffracted up to 3.10 to 3.77 Å resolution (see table S3). As a consequence, the data are 85.6 % complete at 3.19 Å resolution, 81.9% complete at 3.09 Å resolution, but only 22.6% complete in the 3.09 - 2.76 Å resolution shell. Although all available reflections were included in the refinement in order to obtain the most accurate model, the actual resolution of the final thaumatin model is closer to 3.1 Å.

### Applying dynamical scattering correction

As described in (Clabbers et al., 2019), the overestimation of weaker intensities by dynamical scattering can be reduced by the likelihood-based correction made on observed structure factors. This method has yielded good results on strongly diffracting crystals of small molecules and benefitted a relatively strong lysozyme data set. This approach, however, cannot be applied directly to correct data from weakly diffracting protein crystals as the ones in this study.

Electron diffraction data from protein crystals are inherently weak due to the limited electron dose each crystal can tolerate without noticeable damage. A significant fraction of weak intensities is measured as being negative (in our data sets: 12% for thermolysin, 10% for thaumatin, and 6% for insulin) due to counting statistics (Hattne et al., 2016). For these very weak intensities, the calculation of the structure factor is not straightforward and the truncate procedure implemented in CCP4 (Winn et al., 2011) provides a better estimation of the structure factor than just taking the square root of the intensity for positive values and zero for negative ones (French and Wilson, 1978). Refinement tests on the thermolysin electron diffraction data confirmed that using the truncated structure factors allows more reflections to be used in the refinement, leading to a slightly lower gap between R_work_ and R_free_ values and better stereochemistry. However, using these truncated structure factors for dynamical scattering data correction had led to an unrealistic improvement of the refinement statistics, especially in the high-resolution shell, where truncated structure factors derived from negative intensities are most abundant.

In order to avoid any possible bias in the correction introduced by these reflections, we adapted the likelihood-based correction described in (Clabbers et al., 2019) to calculate the correction parameters by comparing I_obs_ and F_calc_^2^ instead of F_obs_ and F_calc_. Such procedure allows negative intensities to be included in the likelihood-based correction and yields a more realistic estimation of dynamical scattering on weak reflections (figure 1). The corrections were calculated with the final model refined against uncorrected data and the corrected data were then used to run 30 successive Refmac runs of 20 cycles each to ensure that both R_work_ and R_free_ reached convergence. Table 1 shows that the correction always improves both R_work_ and R_free_ while the stereochemistry remains essentially identical. The gap between R_work_ and R_free_ is also slightly smaller, except for the thaumatin data. Moreover, the noise level has decreased in the difference Coulomb potential maps. For thermolysin, the corrected map revealed the presence of a solvent molecule that could be either a water molecule, an OH ion or a Cl ion bridging arginine 203 guanidinium group and the Zn ion (figure 2). For comparison, in the 1FJ3 structure, an acetone molecule is observed at this location, with the acetone oxygen atom forming a similar bridge between the guanidinium group and the Zn ion.

**Figure 1.**
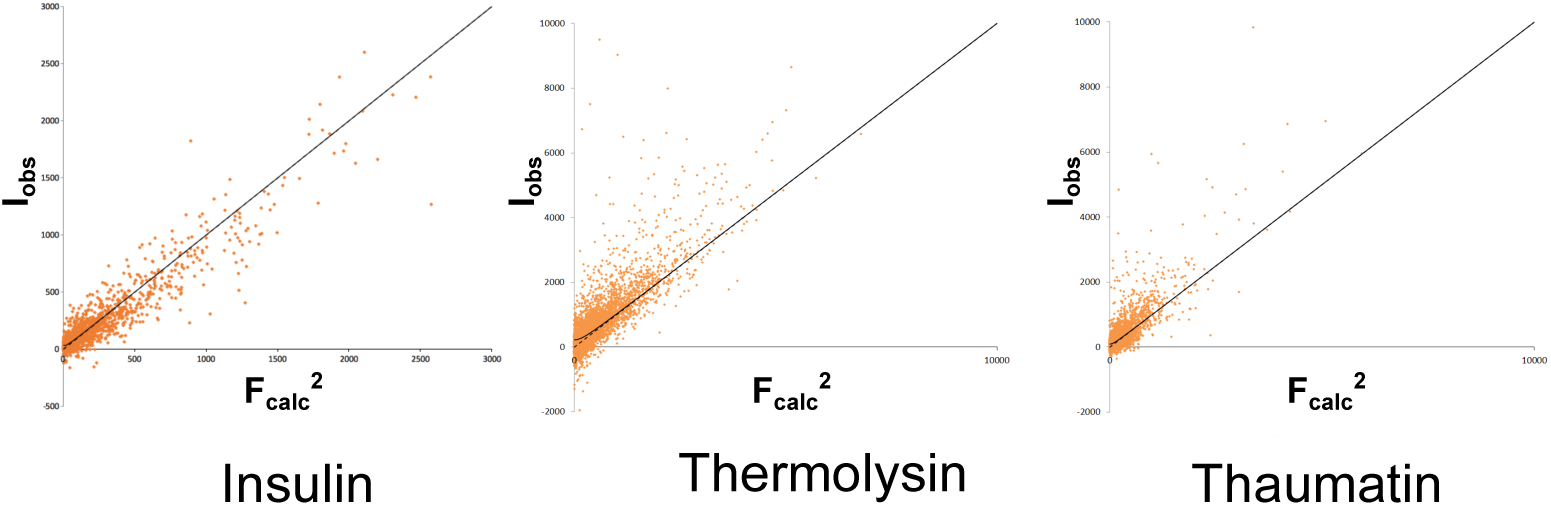
Intensity I_0_ vs 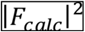 plot for insulin, thermolysin, and thaumatin. Solid line shows the best least squares fit of a hyperbolic function to the observed intensity I_0_. Assuming that 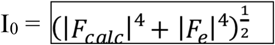, the error term 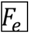 from dynamical scattering can be estimated for the likelihood-based corrections. The dotted line shows the case when I_0_ relates simply to 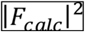.

**Figure 2.**
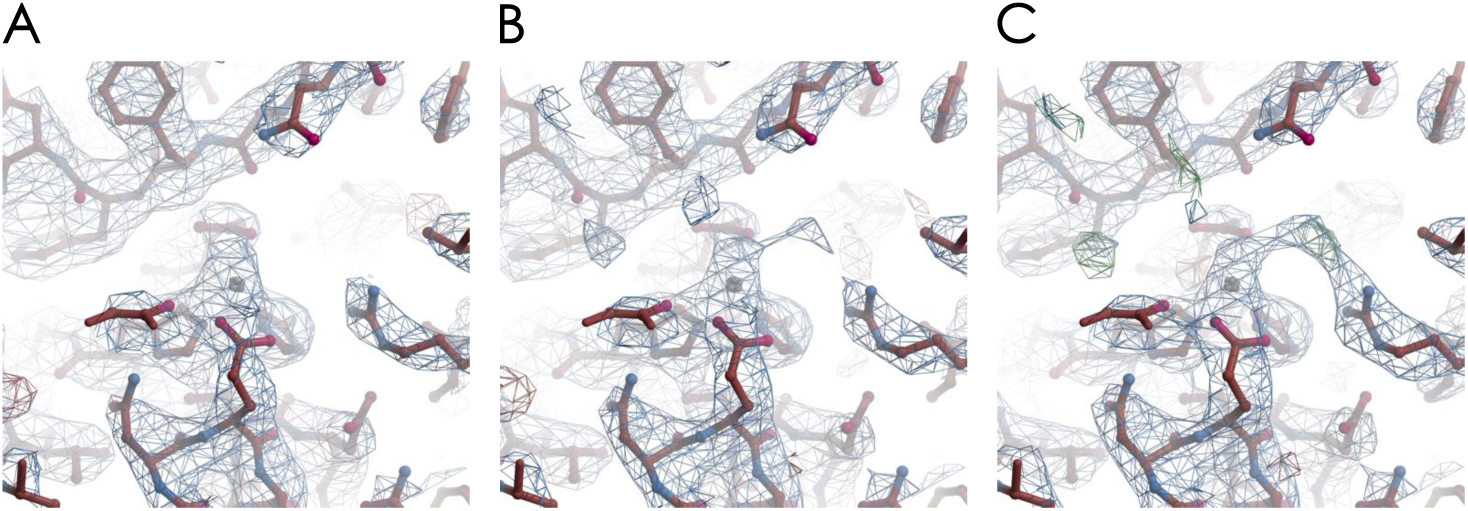
Calculated Coulomb potential map of thermolysin with the atomic model in this study showing the active site with the Zn ion. The structure is determined from data merged from 2 nanocrystals. The {2*F*_obs_ – *F*_calc_} map is contoured at the 1s level and is depicted in blue. The residual {*F*_obs_ – *F*_calc_} map is contoured at the +3s level (depicted in green) and -3s level (depicted in red). (A) Before correcting for dynamical scattering, (B) After correcting for dynamical scattering. (C) After correcting for dynamical scattering and considering the Zn charge. The positive peak in the residual map that appears in the final model could be attributed to a OH or Cl ion, which bridges the Zn ion and the guanidinium group capping the arginine 203.

### Charge analysis of bound ions

Since the electron atomic scattering factor depends strongly on the charge of the atom, we next evaluated the possibility of using the electron diffraction data collected in this study to analyze the charge state of metal ions present in insulin (Zn) and thermolysin (Zn and Ca).

The formal charges of both Zn and Ca ions are +2 in the crystallization solution as the Zn and Ca ions originate from the dilution of ZnCl_2_ and CaCl_2_ salt, respectively. However, according to *ab initio* calculation or electronegativity equalization methods, the actual charge of Zn atoms in protein molecules is often lower and may vary between +0.5 and +1.1, depending on the Zn ligands and the calculation method (Abdallah et al., 2009; Shen et al., 1990). For instance, the Zn ion present in αB-crystallin, which is bound to 3 histidine and a glutamic acid adopting a tetrahedral geometry as in insulin, has been estimated to have a partial charge of +0.72 by electrostatic surface potential calculation (Coi et al., 2005).

In our insulin X-ray structure, a Zn ion (and its symmetry mates) was observed for both chains (B and D) at full occupancy. This Zn ion lies on the crystallographic 3-fold axis and is coordinated by His10 NE2 (Zn-His10 NE2 distance 2.05 Å). The Zn B-factor (28.8 Å^2^ and 27.8 Å^2^) is close to that of His10 NE2 (27.8 Å^2^ and 28.5 Å^2^). The similarity of the B-factors of Zn and His10 suggests a strong binding of the Zn ion. In our refined electron diffraction structure, only one Zn atom of the insulin could be modeled. This Zn atom is chelated to histidine 10D with a similar coordination distance (1.96 Å). When a full occupancy neutral Zn atomic scattering factor is used, the atom has a B-factor of 19.5 Å^2^, which remains essentially the same after refinement against data corrected for multiple scattering (19.6 Å^2^). As observed in table 2, this Zn B-factor is surprisingly lower than the B-factor of the chelating histidine 10D (31 Å^2^). The discrepancy suggests that the use of a positively charged Zn atomic scattering factor is more appropriate for this position.

**Table 2:**
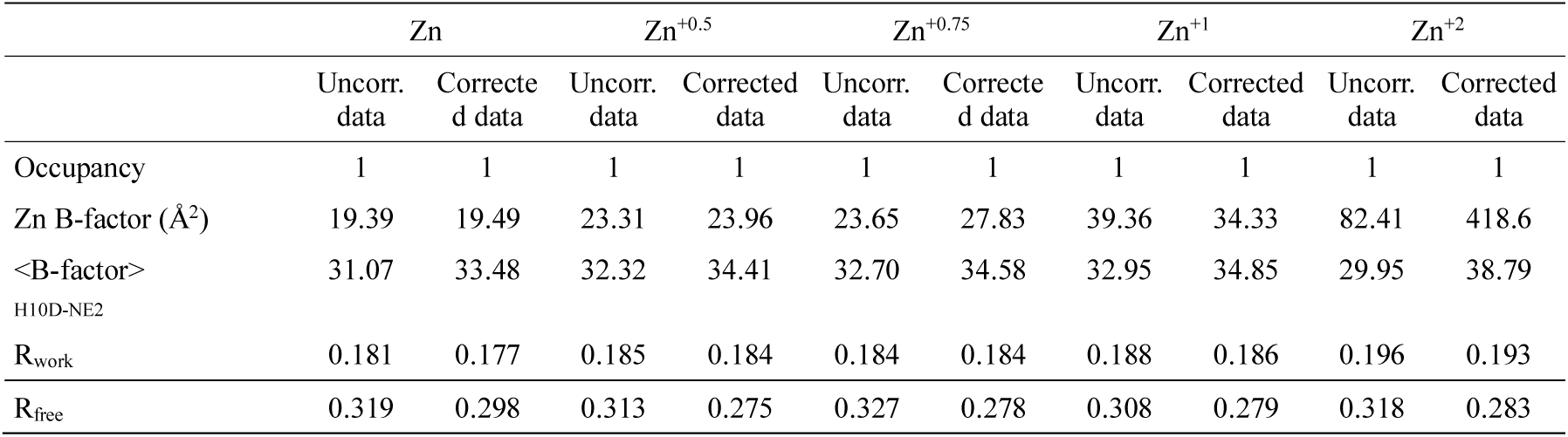
Comparison of refinement statistics on insulin. Uncorrected data and data corrected for dynamical scattering as well as different charge states of the Zn atom are compared.

|  | Zn |  | Zn<sup>+0.5</sup> |  | Zn<sup>+0.75</sup> |  | Zn<sup>+1</sup> |  | Zn<sup>+2</sup> |  |
| --- | --- | --- | --- | --- | --- | --- | --- | --- | --- | --- |
|  | Uncorr.<br>data | Corrected<br>data | Uncorr.<br>data | Corrected<br>data | Uncorr.<br>data | Corrected<br>data | Uncorr.<br>data | Corrected<br>data | Uncorr.<br>data | Corrected<br>data |
| Occupancy | 1 | 1 | 1 | 1 | 1 | 1 | 1 | 1 | 1 | 1 |
| Zn B-factor (Å<sup>2</sup>) | 19.39 | 19.49 | 23.31 | 23.96 | 23.65 | 27.83 | 39.36 | 34.33 | 82.41 | 418.6 |
| <B-factor><br>H10D-NE2 | 31.07 | 33.48 | 32.32 | 34.41 | 32.70 | 34.58 | 32.95 | 34.85 | 29.95 | 38.79 |
| R<sub>work</sub> | 0.181 | 0.177 | 0.185 | 0.184 | 0.184 | 0.184 | 0.188 | 0.186 | 0.196 | 0.193 |
| R<sub>free</sub> | 0.319 | 0.298 | 0.313 | 0.275 | 0.327 | 0.278 | 0.308 | 0.279 | 0.318 | 0.283 |

We analyzed our data following the approach that Yonekura *et al*. employed to analyze electron diffraction data of catalase and β-galactosidase crystals (Yonekura and Maki-Yonekura, 2016). We first calculated the electron atomic scattering factors for different charge states of Zn and Ca ions (+0.5, +0.75, +1.0 and +2.0) in the resolution range of 25 to 1 Å and proceeded to a 5 gaussians parametrization in order to use these atomic scattering factors in Refmac (table s4). We obtained a R_scat_ value, as defined by Yonekura et al. (Yonekura and Maki-Yonekura, 2016), below 0.23 %, indicating that the 5 gaussians parametrization provided an accurate model for the electron atomic scattering factors of Zn and Ca ions (figure 3A). The last refinement step with Refmac was then performed using the different Zn atomic scattering factor parameters. The B-factors and the refinement statistics are shown in table 2. Given that the Zn ion should have a similar B-factor as its chelating histidine (as is the case in the X-ray insulin structure), we estimated the Zn ion charge to be between +0.75 and +1. This estimate is supported by the inspection of the Coulomb potential map for the different charge states, as shown in figure 4, which shows a good agreement between the map and the model for the case of +0.75 Zn charge.

**Figure 3.**
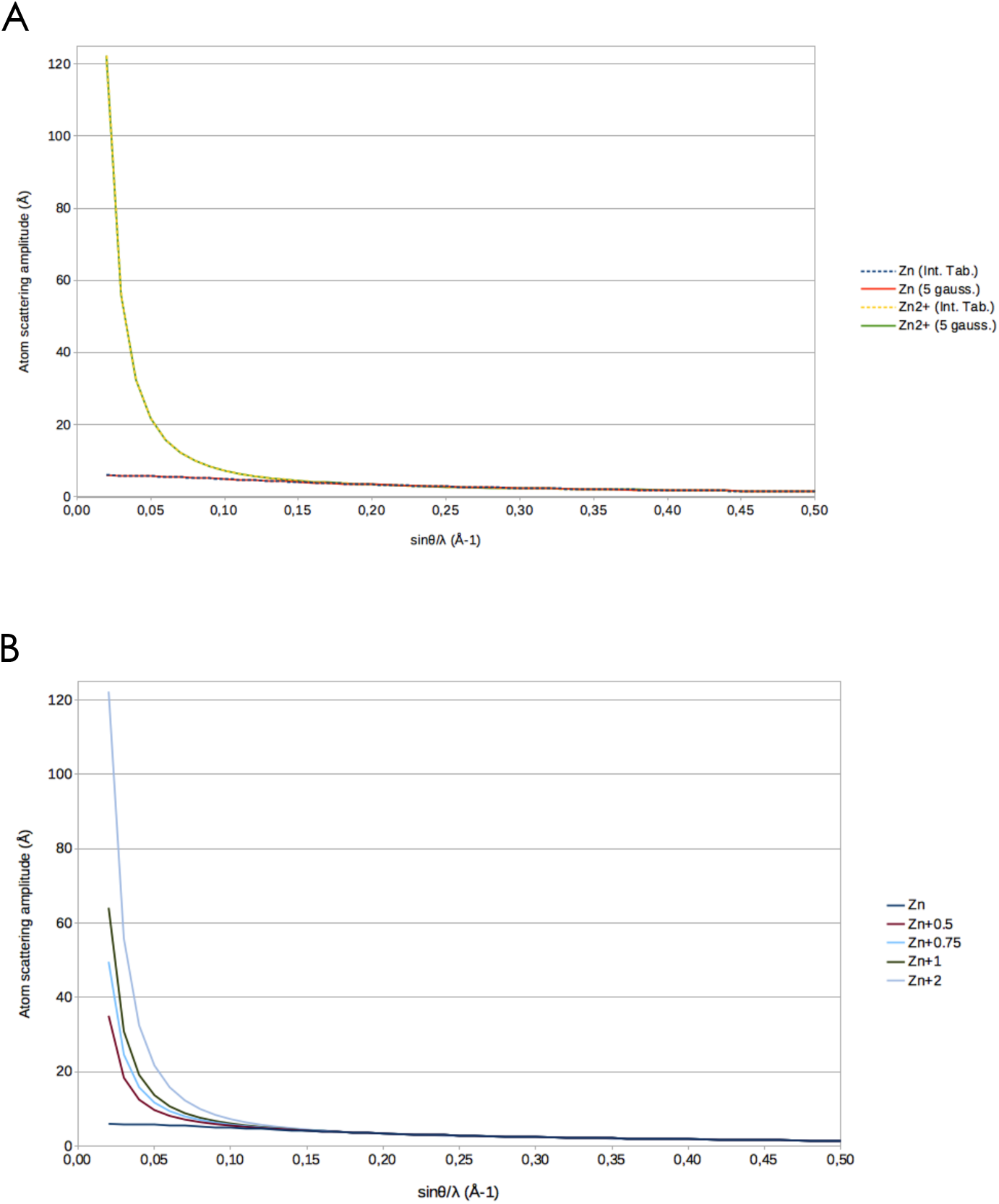
Electron scattering factors for charged Zn atoms. (A) The 5 gaussians parametrizations for neutral and +2 charged Zn atoms follow the red and green solid lines, respectively. The electron atomic scattering curves derived from the International Tables for Crystallography (Vol. C, 2006) for Zn and Zn^2+^ are shown by the blue and yellow dashed lines. The quasi perfect fit illustrates the accuracy of the 5 gaussians parametrization in the 25-1 Å resolution range and the very low R_scat_ values (see table S4). (B) Atomic scattering factors for various charge states of Zn, as calculated from the linear combination of neutral and +2 charge states.

**Figure 4.**
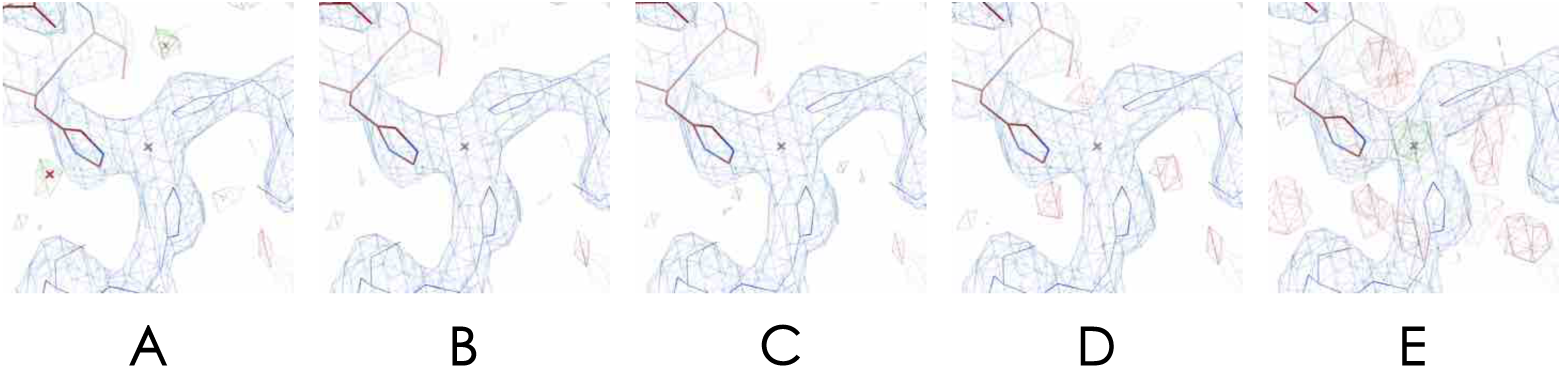
The atomic model and the calculated Coulomb potential map of insulin at one of the Zn ion sites for different assigned Zn charge states. The {2*F*_obs_ – *F*_calc_} map is contoured at the 1σ level and is depicted in blue. The residual {*F*_obs_ – *F*_calc_} map is contoured at the +3σ level (depicted in green) and -3σ level (depicted in red). (A) neutral, (B) +0.5, (C) +0.75, (D) +1.0, (E)+2.0. The Zn ion is coordinated by three histidines. The +0.75 charge yields the optimal residual map.

The same procedure was applied to the thermolysin structure, in which one Zn ion and 4 Ca ions were identified in the Coulomb potential map. Initially, different Zn charge states were tested (0, +0.5, +0.75, +1.0, and +2.0). Unexpectedly, the Zn B-factor obtained with uncorrected data did not smoothly increase with the charge, as would be expected given the increase of the atomic electron scattering factor with the charge value (figure 3B). As shown in table 3, the Zn B-factor for the uncorrected data is relatively stable for charges between 0 and +1 (with even a slight decrease at +1) and increases for charge +2. In contrast, the Zn B-factor obtained with data corrected for dynamical scattering regularly increases with the charge, demonstrating that the data correction has improved the agreement with Wilson statistics and resulted in more realistic B-factors. Comparing the B-factor of Zn and that of the chelating atoms from data corrected for dynamical scattering (table 3) shows that the actual Zn charge is close to +0.75 for thermolysin, as in the case of insulin.

**Table 3:** Comparison of refinement statistics on Thermolysin. Uncorrected data and data corrected for dynamical scattering as well as different charge states of the Zn atom are compared.

|  | Zn |  | Zn<sup>+0.5</sup> |  | Zn<sup>+0.75</sup> |  | Zn<sup>+1</sup> |  | Zn<sup>+2</sup> |  |
| --- | --- | --- | --- | --- | --- | --- | --- | --- | --- | --- |
|  | Uncorr.<br>data | Corrected<br>data | Uncorr.<br>data | Corrected<br>data | Uncorr.<br>data | Corrected<br>data | Uncorr.<br>data | Corrected<br>data | Uncorr.<br>data | Corrected<br>data |
| Zn B-factor ($\text{\AA}^2$) | 16.34 | 24.25 | 16.07 | 28.23 | 17.47 | 35.78 | 15.35 | 43.38 | 37.11 | 84.27 |
| <B-factor><br>H142-NE2, H146-NE2, E143-OE2,<br>E166-OE2, Y157-OH, H231-NE2 | 28.51 | 33.92 | 28.81 | 32.50 | 29.10 | 33.01 | 28.74 | 33.59 | 28.51 | 32.33 |
| R<sub>work</sub> | 0.210 | 0.184 | 0.211 | 0.175 | 0.211 | 0.175 | 0.211 | 0.175 | 0.202 | 0.179 |
| R<sub>free</sub> | 0.292 | 0.238 | 0.292 | 0.240 | 0.292 | 0.240 | 0.298 | 0.240 | 0.298 | 0.251 |

The charge of the 4 Ca ions in thermolysin was investigated using the same approach and the results are shown in table 4. The charge of the Zn ion was set to +2 for the uncorrected data and +0.75 for the corrected data because these charge values had yielded the most realistic Zn B-factor compared to the B-factor of its ligands, respectively. Similar to the case of the Zn ion, the uncorrected data led to an unexpected evolution of the B-factors whereas the corrected data exhibited a smooth increase of Ca B-factors with the charge value. Examining the entries in table 4 suggests that the actual charge of the Ca ions is between +0.75 and +1. The actual value is likely to be closer to +1, where the Ca B-factor is higher or comparable to all the chelating residues B-factors. It is interesting to note that the R_work_ and R_free_ values are not much affected by the charge of either Zn or Ca ions, for charges in the 0 to +1 range. Therefore, these indicators cannot be reliably used to estimate the actual atomic charge. However, R_work_ and R_free_ are systematically higher significantly when the +2 formal charge is used, further supporting our analysis based on the B-factor.

**Table 4:** Comparison of refinement statistics on Thermolysin. Uncorrected data and data corrected for dynamical scattering as well as different charge states of the Ca atoms are compared. The charge of Zn was set to +2 for the uncorrected data and +0.75 for the corrected data, as they provided the most realistic B-factor value for the Zn atom.

|  | Ca |  | Ca<sup>+0.5</sup> |  | Ca<sup>+0.75</sup> |  | Ca<sup>+1</sup> |  | Ca<sup>+2</sup> |  |
| --- | --- | --- | --- | --- | --- | --- | --- | --- | --- | --- |
|  | Uncorr.<br>data | Corrected<br>data | Uncorr.<br>data | Corrected<br>data | Uncorr.<br>data | Corrected<br>data | Uncorr.<br>data | Corrected<br>data | Uncorr.<br>data | Corrected<br>data |
| Ca1 B-factor (Å<sup>2</sup>) | 18.21 | 28.83 | 15.08 | 33.47 | 14.87 | 36.22 | 16.29 | 38.29 | 12.09 | 61.91 |
| <B-factor><br>D138-OD2, E177-OE1, E177-OE2,<br>D185-OD1, E187-O, E190-OE1,<br>E190-OE2 | 35.09 | 38.39 | 34.58 | 40.36 | 33.53 | 41.59 | 32.68 | 38.42 | 28.02 | 36.57 |
| Ca2 B-factor (Å<sup>2</sup>) | 29.56 | 38.61 | 34.39 | 43.84 | 34.92 | 46.92 | 30.61 | 56.51 | 62.45 | 103.78 |
| <B-factor><br>E177-OE2, N183-O, D185-OD1,<br>E190-OE2 | 36.58 | 39.84 | 36.77 | 41.61 | 36.20 | 42.69 | 34.93 | 40.43 | 32.32 | 39.55 |
| Ca3 B-factor (Å<sup>2</sup>) | 18.51 | 26.38 | 21.98 | 26.01 | 18.27 | 26.73 | 11.63 | 30.88 | 28.80 | 52.69 |
| <B-factor><br>D57-OD1, D57-OD2, D59-OD1,<br>Q61-O | 25.70 | 30.86 | 24.18 | 30.85 | 22.28 | 31.64 | 21.17 | 28.54 | 17.98 | 27.40 |
| Ca4 B-factor (Å<sup>2</sup>) | 31.50 | 28.46 | 39.51 | 32.15 | 34.62 | 34.06 | 24.33 | 37.59 | 46.14 | 58.75 |
| <B-factor><br>E190-O, Y193-O, T194-O, T194-OG1, I197-O, D200-OD1 | 39.42 | 43.74 | 37.09 | 44.56 | 35.72 | 45.34 | 35.47 | 41.85 | 28.89 | 41.56 |
| R<sub>work</sub> | 0.202 | 0.175 | 0.200 | 0.179 | 0.201 | 0.183 | 0.205 | 0.184 | 0.212 | 0.201 |
| R<sub>free</sub> | 0.298 | 0.240 | 0.309 | 0.243 | 0.303 | 0.246 | 0.298 | 0.251 | 0.313 | 0.268 |

We also noticed that the Coulomb potential map is rather weak around the various carboxylic groups involved in the coordination of the Zn or Ca ions. Accordingly, we tested applying a negative charge to the oxygen atom of these carboxylic groups. Electron atomic scattering factors for this partially charged oxygen atom were taken from Yonekura et al. (Yonekura and Maki-Yonekura, 2016). Charges from -0.1 to -0.5 were tested and a charge of -0.3 has resulted in the lowest R factor.

## Discussion

Our results confirm that a statistical correction for the overestimation of the intensity of weak reflections arising from dynamical scattering can be generally extended to protein crystals. The correction procedure was originally developed for organic molecule electron diffraction data and so far, had only been demonstrated for protein data using a single lysozyme test case. Here, it has been tested on three different protein crystals and in all the cases, it improved both the refinement statistics and the Coulomb potential map when the corrections are calculated from intensities instead of structure factors, a procedure that allows negative intensities from weak reflections to be included.

Moreover, the correction for dynamical scattering of the data significantly improves the stability of B-factor refinement of ions, which allowed us to estimate the charges of ions in this study. We were able to analyze the charge of several metal ions in insulin and thermolysin crystals, as it has been done for other protein-metal complexes (Yonekura et al., 2015). The Zn ion charge derived from our electron diffraction data agrees well with the partial charge estimated by the electrostatic surface potential calculation for a Zn ion in a similar binding site (Coi et al., 2005). Ca ions in thermolysin are estimated to have a partial charge close to +1.

At first sight, the quality of our electron diffraction data seems inferior compared to X-ray diffraction data even after corrections, in terms of both resolution and I/σ(I) ratio. The thermolysin and thaumatin crystals originate from the same batch of nanocrystals previously analyzed by serial crystallography using an X-FEL source, which yielded structures with a resolution of 2.1 Å for both thermolysin (Hattne et al., 2014) and thaumatin (Nass et al., 2016). However, after the X-FEL experiments, the protein had recrystallized into larger crystals with a size range of 10-50 µm, which is orders of magnitude too large for electron diffraction experiments. We therefore had to crush these crystals by vortexing the crystals with Teflon beads (see Materials and Methods). We cannot exclude the possibility that this compromised the internal ordering of the crystals.

Nonetheless, we believe that the inferior quality of the present electron diffraction data can be largely explained by the difference in the number of unit cells contributing to the signals in the two types of experiments. The electron diffraction results came from only a few thin crystals. These crystals had dimensions in the sub-micron range with a maximal thickness of up to ∼200 nm. In comparison, about 11600 and 125000 crystals of average size 2×3×1 µm^3^ and 3×3×5 µm^3^ contributed to the X-FEL diffraction data of thermolysin and thaumatin, respectively. Given the size of the X-FEL X-ray beam (2.25 µm^2^), the diffracting volume per crystal are in the 2-7 µm^3^ and 7-11 µm^3^ ranges, for thermolysin and thaumatin, respectively. There are thus around six to seven orders of magnitude difference in crystal volume used in the two types of experiments. This difference may fully account for the difference in resolution limit. Similar differences in crystal volume can also explain the higher quality X-ray synchrotron data for the insulin crystals over the electron diffraction data.

In order to fill this gap in data quality, serial electron crystallography is developing quickly and yields a substantial improvement in structure quality (Smeets et al., 2018). Very recently, serial electron diffraction data obtained from naturally produced granulovirus polyhedrin crystals (Bücker et al., 2020) led to higher resolution data than a serial crystallography X-FEL experiment using crystals of comparable size (Gati et al., 2017). Such results and our results presented here experimentally confirm the added value of electron diffraction as a structural biology technique, especially in cases when only sub-micron sized crystals are available and when charge states of component atoms or groups are of interest.

## Materials and methods

### Crystallization

Insulin from bovine pancreas (Sigma I1882) was dissolved in double-distilled water to a concentration of 26 mg/ml in a 1.5 ml reagent tube. To improve the dissolution, the tube was gently agitated for approximately one hour at about 38 °C until the solution remained clear. In a separate 1.5 ml reaction tube the insulin solution was mixed with the crystallization solution (50mM MES pH 6.5, 10mM ZnCl2, 10 mM NaCl in double-distilled water) at a 1:4 ratio with a total volume of 250 µl. The tube was vortexed for 30 seconds directly after the two solutions have been added. The crystallization in this batch setup appeared immediately and crystals in the size range of 10 µm down to ∼50 nm could be observed. Thaumatin and thermolysin crystals were a kind gift of Dr Ilme Schlichting and were from the same batch as used for XFEL diffraction experiments (Nass et al., 2016).

### Crystal handling and freezing

Crystals of 10–50 µm in each dimension were diluted in distilled water (insulin and thaumatin) or buffer (thermolysin) and vortexed for a few minutes in an Eppendorf tube with a 3/32” PTFE bead (Smart Parts) for insulin and 2.381 mm PTFE MicroSeed beads (Molecular Dimensions, MD2-14) for thaumatin and thermolysin. Vortexing time was adjusted in order to obtain a solution of crystals that contained a majority of sub-micrometer crystals that could be used for electron diffraction.

The crystals were then applied onto electron microscopy grids and vitrified. For the insulin sample, 2-3 µl of crystal solution were deposited on one side of either Quantifoil® or lacey copper grids. Extra solution was manually blotted away from the side opposite to the sample deposition side of the grid in order to keep the maximum number of crystals on the grid and plunge frozen in liquid ethane with an artisan plunge freezer. Grids with crystals from thermolysin and thaumatin were prepared using a Vitrobot Mark IV (ThermoFisher Scientific) with 100% humidity at 20 °C. A volume of 2-3 µl of the crystal suspension was applied onto glow-discharged, 300 mesh, copper lacey grids. After blotting for 3-4 s, grids were plunge frozen in liquid ethane cooled by liquid nitrogen.

### Electron diffraction data collection and data processing

The electron diffraction data were collected on a Polara and a Talos cryo-electron microscope (Thermo Fisher) both operated at 200 keV and equipped with a hybrid pixel Timepix detector. Electron diffraction crystallographic data were collected on 7 crystals for insulin (table s1), on 2 crystals for thermolysin (table s2), and on 10 crystals for thaumatin (table s3). All data sets were processed with the XDS package (Kabsch, 2010). The structure of thermolysin and thaumatin were obtained by merging several diffraction data sets of cryo-cooled sub-micron-sized 3D crystals. Two data sets for the thermolysin structure and ten data sets for the thaumatin structure were merged. The two thermolysin crystals were continuously rotated from 17° through 34° with 0.186°/frame and 0.253°/frame in a 2 µm diameter beam. For thaumatin, all ten data sets were continuously rotated starting at 7° and ending at 14°. The data sets of thermolysin and thaumatin showed a decrease in the intensity of reflections with time and the last frames were thus discarded. The data were integrated with XDS and merged with XSCALE to 3.26 Å resolution for thermolysin and 2.76 Å resolution for thaumatin, resulting in an overall completeness of 84% and 66% (table 1). For the insulin data sets, a significant decrease in the intensity of the highest resolution reflections was observed after about 60 s exposure time. Therefore, only the first 50 to 90 frames were used for data processing. Data sets 3 to 7 were merged with XSCALE providing a rather complete 3.25 Å resolution set of unique reflections (table 1). Statistics of data sets 1 and 2 indicated the presence of a merohedral twinning and were thus discarded.

### X-ray data collection and data processing

A 50 µm crystal was taken from the crystal preparation used in electron diffraction experiments, soaked for about 30 s in a cryoprotecting solution containing 25 % of glycerol, and flash frozen in liquid nitrogen. Crystallographic data were collected at 110 K on the BM30A beamline of the European Synchrotron Research Facility (E.S.R.F.), using an ADSC Quantum 315r CCD detector. Data were processed with the XDS package (Kabsch, 2010). A complete 2.3 Å resolution data set was obtained as shown in table 1.

### Parametrization of electron atomic scattering factors for partially charged ions

Electron scattering factors for Zn and Zn^2+^ were taken from the International Table for Crystallography vol. C and were parametrized with a 5 gaussian model as required by Refmac (the fit to determine the 5 gaussian parameters was performed with Fityk, in the range of 25–1 Å). Electron scattering factors for intermediate charges were approximated by a linear combination of Zn and Zn^2+^ atomic scattering factors (Yonekura and Maki-Yonekura, 2016) and parametrized by the same method (table s4). The library of electron atomic scattering factors provided by ccp4 (atomsf_electron.lib) was complemented with the different charge states of Zn and used for refinement in Refmac.

### Structure refinement

The structures were solved by molecular replacement with PHASER from the CCP4 suite of programs (Winn et al., 2011) using the bovine insulin structure from the 2A3G PDB entry, the *Thaumatococcus daniellii* thaumatin structure from the 6C5Y PDB entry and the *Bacillus thermoproteolyticus* thermolysin structure from the 1FJ3 PDB entry as search models. The final structures were obtained by several rounds of manual building with the COOT software (Emsley and Cowtan, 2004) and maximum likelihood refinement with Refmac5 from the CCP4 suite. For the refinement against electron diffraction data, the electron atomic form factors provided by the ccp4 library (atomsf_electron.lib) were used by specifying “source EC” in the Refmac5 script. The ccp4 library was complemented with the different charge states of Zn, Ca and O and used for refinement in Refmac. A likelihood-based correction was performed to reduce the overestimation of lower intensities by dynamical scattering (Clabbers et al., 2019). The refinement statistics are detailed in table 1.

## Acknowledgement

We are very grateful for the support and advice of Igor Nederlof, Martin Weik, and Andrew Leslie. We thank Franck Borel and Jean-Luc Ferrer from BM30A beamline for their help with synchrotron data collection at the European Synchrotron Research Facility (E.S.R.F., Grenoble, France) and Emmanuelle Neumann for her work at the IBS electron microscopy platform. IBS acknowledges integration into the Interdisciplinary Research Institute of Grenoble (IRIG, CEA).

## PDB references

Insulin: 6ZHB (electron diffraction data), 6ZI8 (X-ray data); Thermolysin: 6ZHJ (electron diffraction data); Thaumatin: 6ZHN (electron diffraction data)

---

**Table S1:** Insulin data acquisition and integration statistics of individual crystals used for data merging.

|  | 1 | 2 | 3 | 4 | 5 | 6 | 7 |
| --- | --- | --- | --- | --- | --- | --- | --- |
| **Data collection** | (03) | (05) | (06) | (07) | (Ins-05) | (Ins-20) | (Ins-22) |
| Wavelength (Å) | 0.02508 |  |  |  |  |  |  |
| Frame exposure (s) | 0.6 | 1.5 | 1.2 | 1.2 | 0.6 | 0.6 | 0.6 |
| Number of frames | 50 | 60 | 65 | 60 | 80 | 70 | 90 |
| Total exposure time (s) | 30 | 90 | 78 | 72 | 48 | 42 | 54 |
| $\Phi_{\text{total}}$ (°) | 9.3 | 19.1 | 16.6 | 21.9 | 30.2 | 26.5 | 34.0 |
| $\Delta\phi$ (°/frame) | 0.1855 | 0.3179 | 0.2556 | 0.3654 | 0.3779 | 0.3779 | 0.3779 |
| Detector distance (mm) | 900 | 900 | 900 | 900 | 1890 | 1890 | 1890 |
| **Data processing** |  |  |  |  |  |  |  |
| Space group | *R*3 (H3) | *R*3 (H3) | *R*3 (H3) | *R*3 (H3) | *R*3 (H3) | *R*3 (H3) | *R*3 (H3) |
| Unit cell dimensions |  |  |  |  |  |  |  |
| a, b, c (Å) | a=b=82.54<br>c=33.50 | a=b=82.56<br>c=33.51 | a=b=82.41<br>c=33.15 | a=b=82.40<br>c=33.46 | a=b=82.65<br>c=33.67 | a=b=83.05<br>c=33.64 | a=b=82.46<br>c=34.02 |
| $\alpha, \beta, \gamma$ (°) | 90, 90, 120 | 90, 90, 120 | 90, 90, 120 | 90, 90, 120 | 90, 90, 120 | 90, 90, 120 | 90, 90, 120 |
| Resolution (Å) | 22.6 – 3.53<br>(5.0 – 3.53) | 17.1 – 3.12<br>(3.49 – 3.12) | 23.8 – 3.12<br>(3.49 – 3.12) | 20.60 – 3.02<br>(3.31 – 3.02) | 24.53 – 3.30<br>(3.69 – 3.30) | 21.14 – 3.17<br>(3.54 – 3.17) | 30.73 – 3.67<br>(4.24 – 3.67) |
| $R_{\text{meas}}$ | 0.297 (0.313) | 0.254 (0.645) | 0.231 (0.619) | 0.238 (0.704) | 0.205 (0.882) | 0.196 (1.04) | 0.133 (0.477) |
| $I/\sigma I$ | 3.63 (3.31) | 3.33 (1.54) | 3.55 (1.44) | 3.88 (1.12) | 3.11 (1.03) | 11.2 (3.35) | 4.47 (2.58) |
| Completeness (%) | 14.6 (14.8) | 30.9 (32.1) | 25.2 (24.8) | 34.2 (36.7) | 38.4 (4.1) | 34.0 (33.7) | 40.0 (39.1) |
| Reflections | 273 (183) | 873 (264) | 731 (213) | 1034 (265) | 616 (204) | 599 (205) | 479 (155) |
| Unique reflections | 153 (100) | 470 (139) | 378 (107) | 571 (149) | 493 (153) | 504 (147) | 380 (131) |
| redundancy | 1.78 | 1.86 | 1.93 | 1.81 | 1.25 | 1.19 | 1.26 |
| $R_{\text{sym}}$ | 0.211 (0.222) | 0.181 (0.457) | 0.171 (0.457) | 0.169 (0.499) | 0.147 (0.624) | 0.141 (0.743) | 0.096 (0.337) |
| $CC_{1/2}$ | 0.946 (0.903) | 0.964 (0.804) | 0.970 (0.823) | 0.955 (0.772) | 0.976 (0.632) | 0.994 (0.465) | 0.987 (0.795) |
| **Correlation with X-ray data (resol 50 – 4.0 Å)** |  |  |  |  |  |  |  |
| Indexation 1 | 0.77 | 0.56 | 0.65 | 0.70 | 0.66 | 0.74 | 0.65 |
| Indexation 2 | 0.22 | 0.30 | -0.03 | 0.06 | 0.03 | 0.03 | 0.14 |

**Table S2:** Thermolysin data acquisition and integration statistics of individual crystals used for data merging.

| **Data collection** | 1 | 2 |
| --- | --- | --- |
| Wavelength (Å) | 0.02508 | 0.02508 |
| Frame exposure (s) | 0.6 | 0.8 |
| Number of frames | 90 | 135 |
| Total exposure time (s) | 54 | 108 |
| $\Phi_{\text{total}}$ (°) | 16.74 | 34.14 |
| $\Delta\phi$ (°/frame) | 0.186 | 0.25289 |
| Detector distance (mm) | 1290.2 | 2345 |
| Exposure dose (e.Å<sup>-1</sup>.s<sup>-1</sup>) | 0.0141 | 0.0141 |
| **Data integration** |  |  |
| Space group | *P*6<sub>1</sub>22 | *P*6<sub>1</sub>22 |
| Unit cell dimensions |  |  |
| a, b, c (Å) | 92.00, 92.00, 127.48 | 91.56, 91.56, 127.70 |
| $\alpha$, $\beta$, $\gamma$ (°) | 90, 90, 120 | 90, 90, 120 |
| Resolution (Å) | 14.58-3.26 (3.34-3.26) | 15.56-3.48 (3.57-3.48) |
| R<sub>meas</sub> | 0.472 (1.00) | 0.745 (1.75) |
| I/σI | 2.10 (1.08) | 1.67 (1.01) |
| Completeness (%) | 70.6 (71.6) | 67.6 (68.8) |
| Reflections | 9351 (679) | 9447 (946) |
| Unique reflections | 3800 (282) | 2994 (220) |

**Table S3:** Thaumatin data acquisition and integration statistics of individual crystals used for data merging.

|  | 1 | 2 | 3 | 4 | 5 | 6 | 7 | 8 | 9 | 10 |
| --- | --- | --- | --- | --- | --- | --- | --- | --- | --- | --- |
| **Data collection** | (21) | (33) | (19) | (30) | (27) | (08) | (14) | (16) | (10) | (09) |
| Wavelength (Å) | 0.02508 |  |  |  |  |  |  |  |  |  |
| Frame exposure (s) | 1.2 | 1.2 | 1.2 | 2.4 | 1.2 | 1.2 | 1.2 | 1.2 | 1.2 | 1.2 |
| Number of frames | 100 | 185 | 127 | 50 | 180 | 180 | 190 | 170 | 150 | 100 |
| Total expo-sure time (s) | 120 | 222 | 152.4 | 120 | 216 | 216 | 228 | 204 | 180 | 120 |
| $\Phi_{\text{total}}$ (°) | 7.3 | 13.505 | 9.271 | 7.3 | 13.14 | 13.14 | 13.87 | 12.41 | 10.95 | 7.3 |
| $\Delta\phi$ (°/frame) | 0.073 | 0.073 | 0.073 | 0.146 | 0.073 | 0.073 | 0.073 | 0.073 | 0.073 | 0.073 |
| Detector distance (mm) | 2345 | 2345 | 2345 | 2345 | 2345 | 2345 | 2345 | 2345 | 2345 | 2345 |
| **Data integration** |  |  |  |  |  |  |  |  |  |  |
| Space group | $P4_12_12$ | | | | | | | | | |
| Unit cell dimensions |  |  |  |  |  |  |  |  |  |  |
| a, b, c (Å) | 57.72, 57.72, 149.17 |  |  |  |  |  |  |  |  |  |
| $\alpha, \beta, \gamma$ (°) | 90, 90, 90 | | | | | | | | | |
| Resolution (Å) | 10.69-2.76<br>(2.86-2.76) | 11.69-3.24<br>(3.37-3.24) | 8.83-3.12<br>(3.34-3.12) | 9.81-3.10<br>(3.27-3.10) | 11.74-3.39<br>(3.54-3.39) | 12.26-3.40<br>(3.54-3.40) | 12.01-3.33<br>(3.47-3.33) | 10.66-3.77<br>(4.03-3.77) | 10.49-3.16<br>(3.32-3.16) | 8.630-3.26<br>(3.52-3.26) |
| $R_{\text{meas}}$ | 0.390<br>(0.998) | 0.417<br>(0.948) | 0.577<br>(1.21) | 0.388<br>(1.06) | 0.456<br>(0.954) | 0.524<br>(1.10) | 0.522<br>(1.11) | 0.764<br>(1.08) | 0.484<br>(1.20) | 0.592<br>(1.85) |
| $I/\sigma(I)$ | 2.58<br>(1.10) | 2.20<br>(1.06) | 1.59<br>(1.01) | 1.97<br>(1.02) | 1.79<br>(1.05) | 1.66<br>(1.00) | 2.31<br>(1.09) | 1.54<br>(1.02) | 2.81<br>(1.12) | 2.26<br>(1.08) |
| Completeness (%) | 21.7<br>(21.9) | 29.7<br>(31.5) | 17.0<br>(19.5) | 20.5<br>(24.1) | 31.0<br>(35.0) | 34.7<br>(38.7) | 31.3<br>(35.9) | 29.1<br>(32.8) | 22.4<br>(22.7) | 15.7<br>(17.5) |
| Reflections | 2682<br>(311) | 2834<br>(397) | 2114<br>(489) | 1690<br>(327) | 2374<br>(358) | 2214<br>(308) | 2280<br>(344) | 1542<br>(368) | 2222<br>(376) | 1333<br>(352) |
| Unique reflections | 1517<br>(145) | 1309<br>(151) | 837<br>(166) | 1029<br>(171) | 1201<br>(153) | 1329<br>(154) | 1282<br>(163) | 837<br>(166) | 1064<br>(138) | 681<br>(150) |

**Table S4:** Parametrization of electron atomic scattering factors for Zn and Ca ions, using the 5 gaussians model. Parameters were fitted to data in the range 0.02 ≤ sinθ / λ ≤ 0.5

|  | a1, b1 | a2, b2 | a3, b3 | a4, b4 | a5, b5 | R<sub>scat</sub> (%) |
| --- | --- | --- | --- | --- | --- | --- |
| Zn<sup>+0.5</sup> | 2.91473<br>2.77406 | 2.83557<br>30.9254 | 4.13357<br>187.481 | 63.2726<br>3600.28 | 15.0154<br>885.604 | 0.18 |
| Zn<sup>+0.75</sup> | 2.96853<br>2.86011 | 22.8485<br>919.772 | 3.0442<br>33.842 | 95.0852<br>3649.59 | 6.2312<br>207.334 | 0.20 |
| Zn<sup>+1</sup> | 3.01042<br>2.92881 | 126.93<br>3682.27 | 30.7484<br>942.7 | 8.42183<br>220.231 | 3.2772<br>36.6377 | 0.21 |
| Zn<sup>+2</sup> | 3.12034<br>3.12016 | 62.9056<br>1000.92 | 254.57<br>3764.19 | 17.7729<br>251.618 | 4.53073<br>47.6443 | 0.22 |
| Ca<sup>+0.5</sup> | 2.84711<br>3.8095 | 3.7506<br>34.3279 | 63.0883<br>3527.94 | 5.76723<br>153.973 | 14.6981<br>836.477 | 0.20 |
| Ca<sup>+0.75</sup> | 2.91939<br>3.92932 | 3.9753<br>37.149 | 7.29405<br>179.917 | 22.5015<br>882.291 | 94.8902<br>3595.47 | 0.22 |
| Ca<sup>+1</sup> | 2.97364<br>4.01986 | 126.761<br>3645.44 | 4.13874<br>39.4108 | 30.4422<br>916.795 | 9.10789<br>200.748 | 0.23 |
| Ca<sup>+2</sup> | 3.11407<br>4.25523 | 4.58655<br>46.2978 | 62.7958<br>997.095 | 254.521<br>3758.58 | 17.7083<br>249.738 | 0.23 |

**Figure S1.**
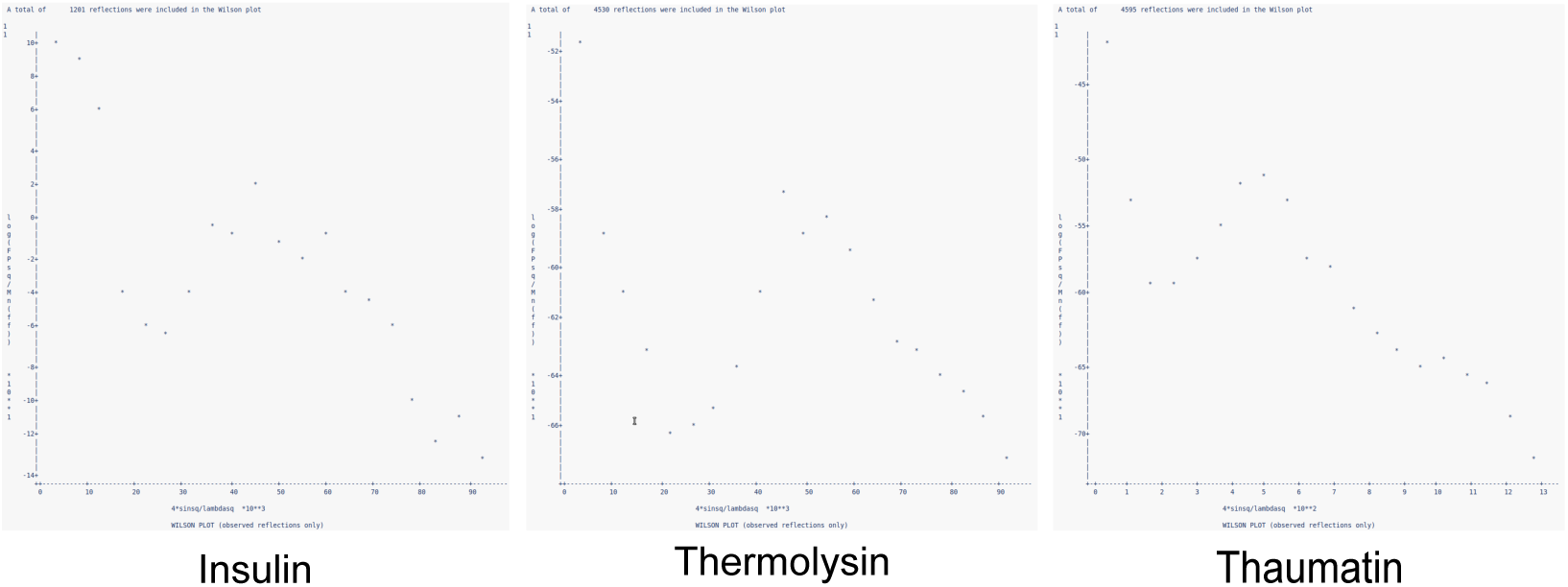
Wilson plots of electron diffraction data for the three proteins.

